# Non-enzymatic assembly of active chimeric ribozymes from aminoacylated RNA oligonucleotides

**DOI:** 10.1101/2021.09.15.460531

**Authors:** Aleksandar Radakovic, Saurja DasGupta, Tom H. Wright, Harry R.M. Aitken, Jack W. Szostak

## Abstract

Aminoacylated tRNAs, which harbor a covalent linkage between amino acids and RNA, are a universally conserved feature of life. Because they are essential substrates for ribosomal translation, aminoacylated oligonucleotides must have been present in the RNA World prior to the evolution of the ribosome. One possibility we are exploring is that the aminoacyl ester linkage served another function before being recruited for ribosomal protein synthesis. The nonenzymatic assembly of ribozymes from short RNA oligomers under realistic conditions remains a key challenge in demonstrating a plausible pathway from prebiotic chemistry to the RNA World. Here, we show that aminoacylated RNAs can undergo template-directed assembly into chimeric amino acid-RNA polymers that are active ribozymes. We demonstrate that such chimeric polymers can retain the enzymatic function of their all-RNA counterparts by generating chimeric hammerhead, RNA ligase, and aminoacyl transferase ribozymes. Amino acids with diverse side chains form linkages that are well tolerated within the RNA backbone, potentially bringing novel functionalities to ribozyme catalysis. Our work suggests that aminoacylation chemistry may have played a role in primordial ribozyme assembly. Increasing the efficiency of this process provides an evolutionary rationale for the emergence of sequence and amino acid specific aminoacyl-RNA synthetase ribozymes, which could then have generated the substrates for ribosomal protein synthesis.

**Significance Statement:** The emergence of the primordial ribosome from the RNA World would have required access to aminoacylated RNA substrates. The spontaneous generation of such substrates without enzymes is inefficient, and it remains unclear how they could be selected for in a prebiotic milieu. In our study we identify a role for aminoacylated RNA in ribozyme assembly, a longstanding problem in the origin of life research. We show that aminoacylated RNAs, but not unmodified RNAs, rapidly assemble into chimeric amino acid-bridged ribozymes that retain their native enzymatic activity. Our work therefore addresses two key challenges within the origin-of-life field: we demonstrate assembly of functional ribozymes and we identify a potential evolutionary benefit for RNA aminoacylation that is independent of coded peptide translation.

## Introduction

The evolution of ribosomal protein synthesis would have required the presence of aminoacylated RNAs, which are the universal substrates for protein synthesis. Without specialized aminoacyl-RNA synthetase enzymes, the initial aminoacylation of RNA must have been mediated by spontaneous chemical processes. However, current chemical pathways leading to RNA-aminoacylation are inefficient or rely on unstable, activated amino acids. For example, nonenzymatic aminoacylation of the *cis* diol of RNA oligonucleotides with imidazole-activated amino acids affords low levels of aminoacylation,(1, 2) primarily due to the pronounced hydrolytic instability of protonated aminoacyl esters.(3, 4) Alternatively, interstrand transfer of an amino acid from a 5′ phosphate mixed anhydride to the *cis* diol generates aminoacylated oligonucleotides but depends upon the efficient synthesis of phospho-carboxy anhydrides, which are prone to hydrolysis and CO_2_-mediated degradation.(5, 6) Given these limitations, spontaneous synthesis alone may not have been generated sufficient aminoacylated-RNA substrates to facilitate the emergence of a primitive ribosome. Furthermore, aminoacylated-RNAs generated by spontaneous synthesis are likely to have exhibited low sequence and amino acid specificity, which would have made the evolution of coded peptide synthesis problematic. Here we consider a scenario in which the initial, inefficient and minimally specific aminoacylation of RNA played an alternative role that was nonetheless advantageous to primordial protocells. Such a role might have provided a selective pressure favoring the evolution of ribozymes that generated and maintained high levels of RNA aminoacylation. Accordingly, we have sought to identify a role for aminoacylated RNAs that could have been beneficial to primitive protocells, independent of ribosomal protein synthesis.

Early life is thought to have used RNA enzymes, or ribozymes, to catalyze primordial metabolic reactions as well as RNA replication, prior to the evolution of protein enzymes (i.e. the RNA World).(7, 8) In this scenario, the first ribozymes were generated through the nonenzymatic ligation of short RNA oligonucleotides and/or the template directed polymerization of activated nucleotide monomers. While the activation of ribonucleotides with imidazole derivatives has facilitated the assembly and templated copying of short RNAs,(9–13) oligomers long enough to exhibit enzymatic properties have remained out of reach. A major obstacle to efficient ribozyme assembly is the relatively poor nucleophilicity of the terminal *cis* diol of RNA, mandating high Mg^2+^concentrations that are not compatible with fatty acid-based vesicle models of protocells.(14) Carbodiimide activation of phosphate(15, 16) or replacement of the 3′ hydroxyl with an amino group allow for efficient ligation of RNA oligonucleotides,(17–22) but the prebiotic relevance of these approaches remains unclear.

In the 1970s, Shim and Orgel demonstrated that glycylated nucleotides can take part in template-directed polymerization with imidazole-activated nucleotides, forming phosphoramidate linkages between the glycine amine and the 5′-phosphate of an adjacent nucleotide.(23) Building on this discovery, recent investigations have focused on the likelihood of an RNA-amino acid copolymer world(24) and the conditions that permit conjugation of peptido RNAs.(25) The Richert group has recently expanded our understanding of peptidoyl (peptide bridged) RNAs and how they may have played a role in template-directed peptide condensation.(25, 26) However, to our knowledge, chimeric, amino acid-bridged ribozymes created with the building blocks of both proteins and RNA have not yet been reported. In our previous work, short aminoacylated RNAs were shown to form amino acid bridges in non-enzymatic ligation reactions with imidazole-activated oligonucleotides, at rates much higher than observed with unmodified RNA.(27) Here, we leverage this enhanced ligation reactivity to generate three ribozymes with chimeric backbones that perform RNA cleavage, RNA ligation, and RNA aminoacylation, all thought to have been important enzymatic functions in the RNA World. Importantly, we generate the chimeric ribozymes at 2.5 mM Mg^2+^, conditions that are compatible with fatty acid-based vesicles(14) and limit RNA and activated RNA degradation,(28) but generally yield no RNA ligation. Our work reveals a link between aminoacylation chemistry and the non-enzymatic assembly of RNA oligonucleotides into functional ribozymes. This functional link provides a potential rationale for the evolution of ribozyme-catalyzed RNA aminoacylation prior to the evolution of ribosomal protein synthesis.

## Results

Having previously demonstrated the enhanced template-directed ligation of aminoacylated RNAs,(27) we asked whether this strategy could be employed to assemble catalytically active ribozymes containing multiple interspersed amino acid bridges. To explore the functionality of these chimeric ribozymes, we first targeted the assembly of a 37 nt long hammerhead ribozyme from three oligonucleotides via templated ligation. Because efficient prebiotic pathways for RNA aminoacylation have not yet been elucidated, we utilized an aminoacylating ribozyme (Flexizyme)(29) to generate aminoacylated RNA building blocks for ligation. Oligonucleotide substrates for Flexizyme aminoacylation must terminate with 5′-NNCA-3′. To accommodate this requirement, we mutated a wild-type hammerhead ribozyme sequence to introduce two CA dinucleotides at positions that would allow us to assemble the complete ribozyme from three oligonucleotides, with the first two oligonucleotides modified to contain the required 3′ termini for aminoacylation (Figure S1). Upon activation with 2-methylimidazole followed by aminoacylation with glycine and addition of an RNA template, we observed rapid ligation of the glycylated oligonucleotides to yield the full-length 37 nt chimeric hammerhead, which was detectable as early as 5 minutes (Figure 1). In contrast, non-aminoacylated RNA oligomers yielded only trace amounts of full-length product after 24 h. While this result suggested that multiple ligation reactions with aminoacylated RNA could be leveraged to assemble chimeric ribozymes, the formation of a stable duplex between the RNA template and the ribozyme RNA prevented the quantitative measurement of ribozyme activity.

**Figure 1.**
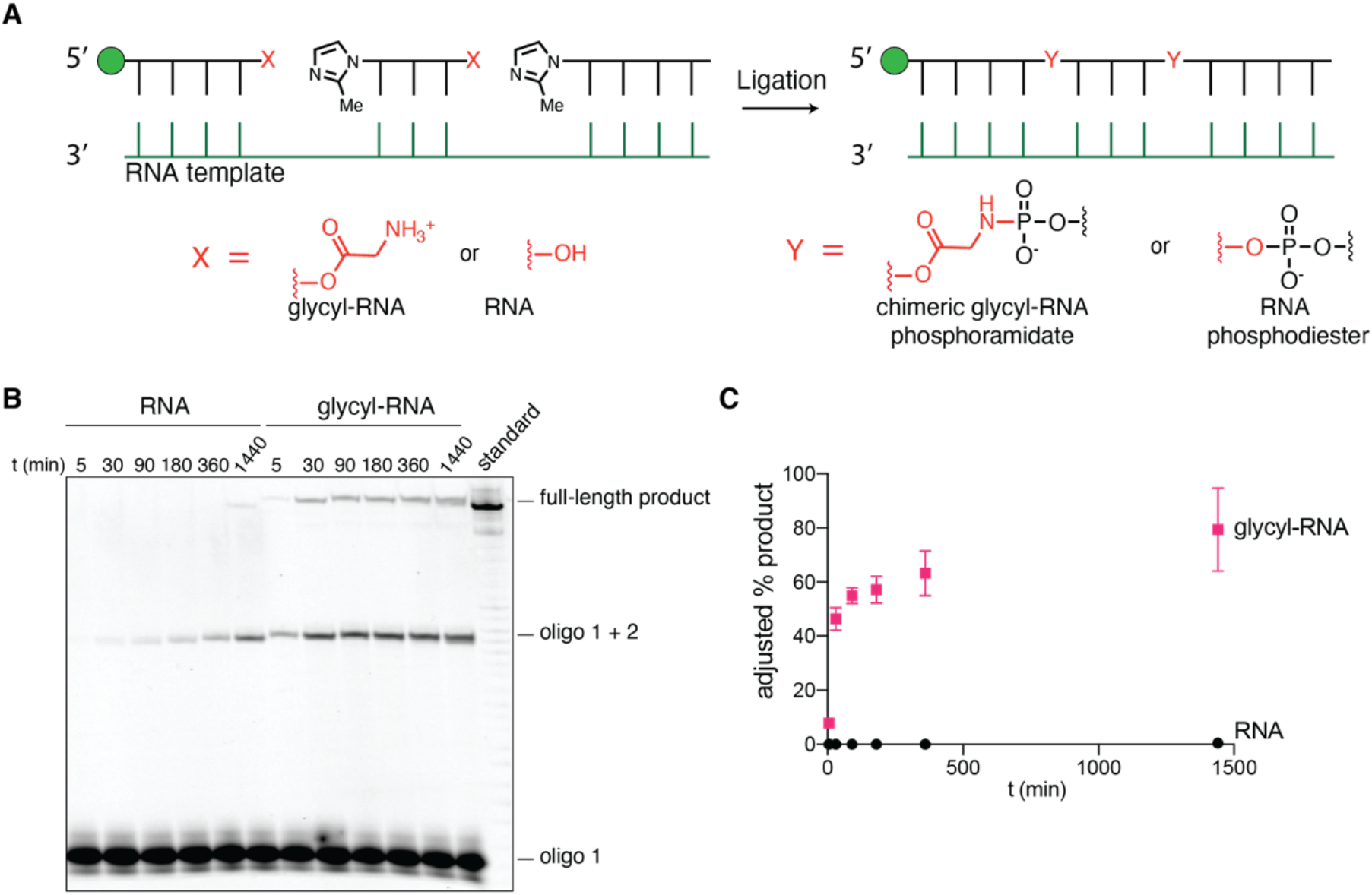
Glycylated RNA oligonucleotides undergo rapid ligation reactions to produce long chimeric polymers. **A** Schematic of the template-directed ligation reaction. **B** Denaturing urea-PAGE of the ligation reactions for non-aminoacylated RNA oligonucleotides (RNA) and glycylated oligonucleotides (glycyl-RNA). Oligo 1 is the starting oligonucleotide, oligo 1 + 2 is the product of the first ligation, and the full-length product is the production of two ligations. **C** Plot of full-length product yields vs. time. Glycylated RNA is represented by pink squares, RNA is represented by black circles. Glycylation yields for oligonucleotides 1 and 2 were measured by acidic urea-PAGE independently, and the adjusted yield was obtained by dividing the raw ligation yield by the aminoacylation percent yield. Reactions were performed in triplicate in 200 mM Na+-HEPES pH 8.0 and 2.5 mM MgCl_2_, with 1.25 *μ*M template and 1.25 μM each oligonucleotide.

To assess the activity of the chimeric hammerhead ribozyme, we modified the assembly process by using a DNA, rather than an RNA, template complementary to the full ribozyme sequence. We have previously found that DNase digestion of similar templates can be used to release pyrophosphate-linked(30) and amino acid-bridged(27) RNAs, and we reasoned that the same strategy would release the full-length chimeric ribozyme, allowing it to fold into the active conformation and bind its substrate (Figure S1C). Although not prebiotically relevant, this approach permitted a systematic assessment of the activity of chimeric ribozymes. Using this method, we assembled four chimeric hammerhead ribozymes using four different amino acids with diverse side chains (Figure 2A, S3). All four chimeric ribozymes cleaved the native hammerhead substrate under single-turnover conditions, albeit at rates that were 20 to 80-fold lower than the all-RNA hammerhead ribozyme (Figure 2B, C). The L-lys linked chimeric hammerhead exhibited the fastest cleavage of the four chimeric ribozymes, demonstrating that bulky, positively charged amino acid side chains can be tolerated within the phosphodiester backbone.

**Figure 2.**
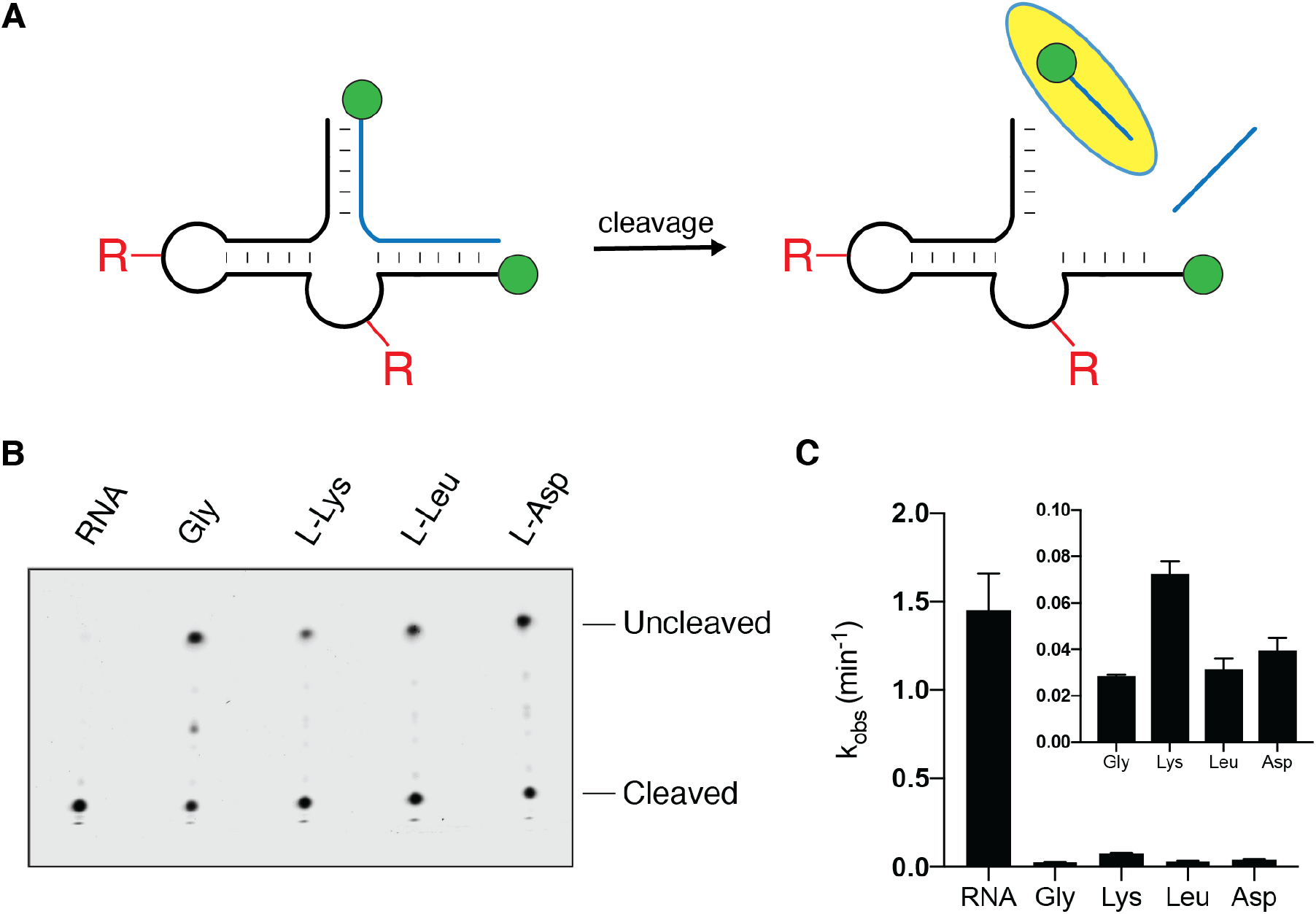
Assembly of active chimeric hammerhead ribozymes with four different bridging amino acids. **A** Diagram of the Hammerhead cleavage reaction. Red R groups represent the four different amino acid bridges. The highlighted oligonucleotide is the cleaved substrate observed in the following panels. **B** Urea-PAGE of the Hammerhead cleavage reactions. RNA is all-RNA ribozyme, while the chimeric ribozymes assembled with different amino acids are abbreviated with the amino acid used in their assembly. Top band: uncleaved fluorescein-labeled Hammerhead substrate; Bottom band: cleaved substrate. **C** Kinetic analysis of the Hammerhead cleavage reactions. The inset shows the observed rate constants for the chimeric ribozymes in the same order as in the main panel (note the distinct y-axis scale). Reactions were performed in triplicate with 0.12 *μ*M ribozyme, 0.1 *μ*M FAM-labeled substrate in the presence of 100 mM Tris-Cl (pH 8) and 3 mM MgCl_2_.

Having shown that amino acid bridged RNAs can exhibit catalytic phosphodiester bond cleavage, we set out to design and assemble a chimeric ribozyme that can catalyze phosphodiester bond formation. As a starting point we chose an *in vitro* selected ligase ribozyme that catalyzes the ligation of imidazole-activated oligonucleotides.(31) First, we tested truncations of the original 70 nt sequence and were able to eliminate 4 base pairs from the first stem, resulting in a 62 nt ribozyme (Figure S4). To accommodate Flexizyme-catalyzed aminoacylation of individual oligomeric building blocks, we screened variants of this truncated sequence containing CA dinucleotides in positions that would enable us to assemble the entire sequence from shorter 3′-aminoacylated, 5′-(2-methylimidazole) activated RNA oligonucleotides (Figure S4). While some variants lost ligase activity, we were able to identify a suitable ribozyme sequence that could be assembled from four oligonucleotides ranging in length from 11 to 18 nts.

We assembled the chimeric ligase ribozyme using RNA oligonucleotides aminoacylated with L-lys because aminoacyl-RNA ligation with this amino acid resulted in the highest yields (Figure 3A, S5).(27) Our choice of L-lys was also informed by the observation that L-lys amino acid bridges were the most well tolerated within the hammerhead ribozyme (Figure 2 B, C). Despite containing three L-lys bridges in its backbone, the chimeric ligase ribozyme was functional, exhibiting half the activity of the corresponding all-RNA ribozyme (Figure 3B, C). Due to the faster hydrolysis of aminoacyl ester-phosphoramidate linkages compared to phosphodiester linkages, we observed a gradual decrease in the amount of full-length ligated product after 60 minutes (Figure 3C). A similarly assembled glycine-bridged ligase was also functional; however, due to the lower yield of full-length product with glycine aminoacylated oligonucleotides we could not quantitate its activity (Figure S6).

**Figure 3.**
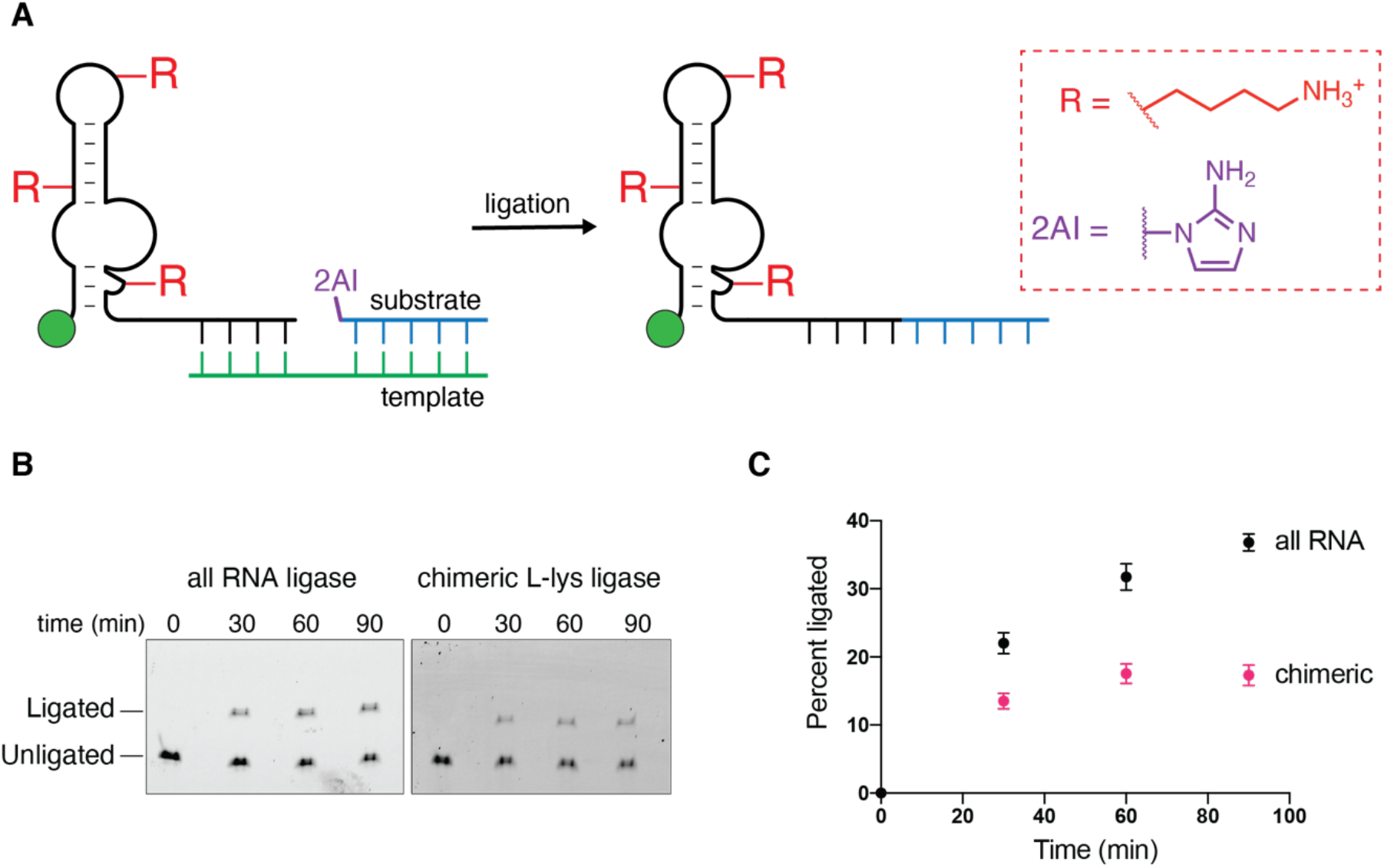
Active chimeric RNA ligase ribozyme assembled with L-lysylated oligonucleotides. **A** Schematic of the RNA ligase reaction. The red R groups represent L-lysine linkages. **B** Urea-PAGE analysis of a representative time-course of the ribozyme-catalyzed ligation reaction. Bottom band: fluorescein-labeled ligase ribozyme; Top band: ribozyme ligated to its substrate. **C** Plot of the time-course of the ligation reaction. Black circles represent the percent of ligated product for the all-RNA ribozyme; gray circles represent the percent of ligated product for the chimeric L-lys ribozyme. Reactions were performed in triplicate with 0.1 *μ*M ribozyme, 0.12 *μ*M template, and 0.2 *μ*M substrate in the presence of 100 mM Tris-Cl (pH 8) and 10 mM MgCl_2_.

The ligation of aminoacylated oligonucleotides to generate a chimeric aminoacyl-RNA synthetase ribozyme could in principle result in an autocatalytic system in which the aminoacylating ribozyme synthesizes its own precursors, which then spontaneously assemble into more of the aminoacylating ribozyme. Even an inefficient initial chemical aminoacylation could potentially be sufficient to initiate such a self-amplifying cascade. As a first step toward such a cascade, we assembled a chimeric aminoacylating ribozyme (Flexizyme) from gly and L-lys aminoacylated oligonucleotides. The wild type Flexizyme sequence contains two CA sequences at positions that allow the assembly of the complete ribozyme from three oligonucleotides (Figure S7). Although both CA sequences are in the catalytic center of the ribozyme, we found that both gly and L-lys bridged Flexizymes were functional (Figure 4, S7, S8). Mutating the Flexizyme to reposition the CA sequences and gly bridges outside the catalytic center did not improve the activity of the chimeric ribozyme (Figure S7). Given that we initially used the all-RNA Flexizyme to aminoacylate the oligonucleotides that were then assembled into the chimeric Flexizyme, it seemed conceivable that contamination of the purified chimeric Flexizyme with traces of the all-RNA Flexizyme could have contributed to the observed aminoacylating activity (see Methods for details on aminoacylated oligonucleotide gel purification). To rule out this possibility, we pretreated the chimeric Flexizymes at pH 12 for 30 s, conditions that hydrolyze aminoacyl esters but not phosphodiesters, prior to the aminoacylation reaction. The chimeric Flexizymes lost all aminoacylating activity after such pretreatment, while the all-RNA Flexizyme retained most of its activity (Figure 4B, C, S8). Similarly to the hammerhead and ligase ribozymes, the L-lys bridged chimeric Flexizyme was the most active among those tested (Figure 4, S8). The L-lys bridged ribozyme aminoacylated 10 % of its RNA substrate after 22 h, demonstrating that enzymatic aminoacylation can emerge from the nonenzymatic ligation of aminoacylated oligonucleotides.

**Figure 4.**
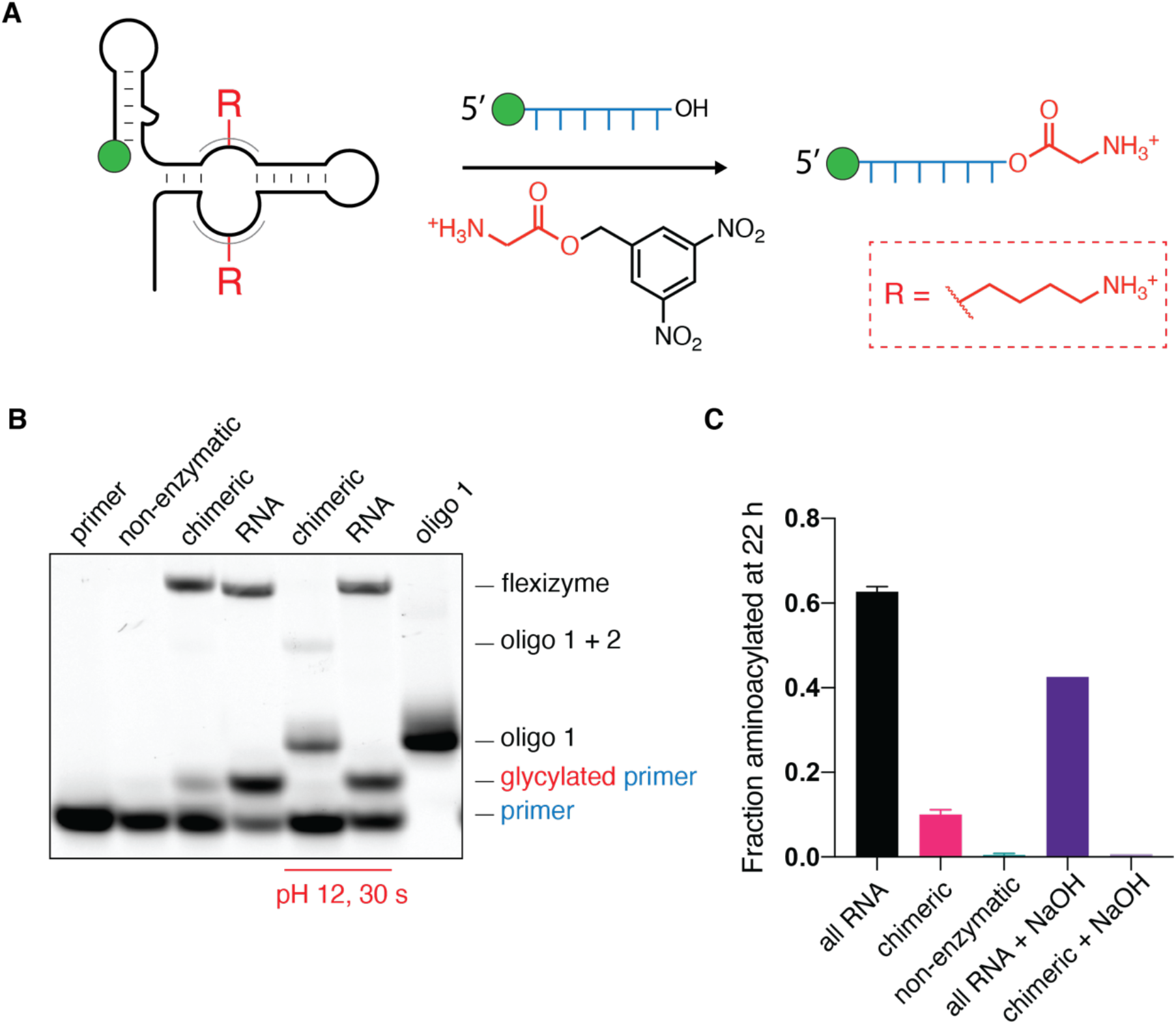
Active chimeric Flexizyme assembled from L-lysylated oligonucleotides. **A** Schematic of the chimeric Flexizyme-catalyzed aminoacylation reaction. Red R groups represent L-lysine linkages. **B** Acidic urea-PAGE analysis of the aminoacylation reaction. Non-enzymatic lane is the background aminoacylation reaction in the absence of any ribozyme. Bottom band: primer to be glycylated; above it is the glycylated primer. Oligonucleotide 1 and oligonucleotide 1 + 2 represent the disassembled products of the chimeric Flexizyme upon aminoacyl ester hydrolysis. **C** Bar plot of the final aminoacylation yields after 22 h (see Figure S8 for full time-course). Reactions were performed in triplicate, except for the alkali-treated reactions, in 50 mM Na+-HEPES pH 8.0 and 10 mM MgCl_2_, with 2.5 *μ*M Flexizyme, 10 *μ*M primer, and 25 mM glycine-DBE.

The chimeric ribozyme assembly method described above involves the use of a DNA template followed by DNase digestion to liberate the product ribozyme. Because neither the long DNA template nor the enzymatic digestion are prebiotically relevant, we sought to assemble a chimeric ribozyme using the splint-assisted assembly strategy (Figure 5A, S9). Splint-assisted assembly has been shown to obviate the need for long template strands(32) and to overcome the inhibition of the assembled ribozyme by the strong binding of the template strand to the ribozyme.(22) Using DNA splints or RNA splints with at least one G*U wobble base-pair led to the formation of the full-length gly bridged hammerhead variant from glycylated oligonucleotides. RNA oligonucleotides lacking the glycyl group failed to generate even trace amounts of full-length product under the same conditions (Figure 5B). To facilitate splint release and substrate binding, we added the hammerhead substrate and heated the reaction at 95 °C for 2 minutes before cooling it to the reaction temperature. Performing the subsequent cleavage reaction at 42 °C, conditions that release the ribozyme from the splints but leave substrate binding unaffected, we observed the expected cleavage of the substrate (Figure S10). Importantly, performing the reaction at 25 °C, conditions that allow both splint and substrate binding, resulted in similar levels of substrate cleavage, indicating that the chimeric hammerhead is functional despite the presence of splints (Figure 5C).

**Figure 5.**
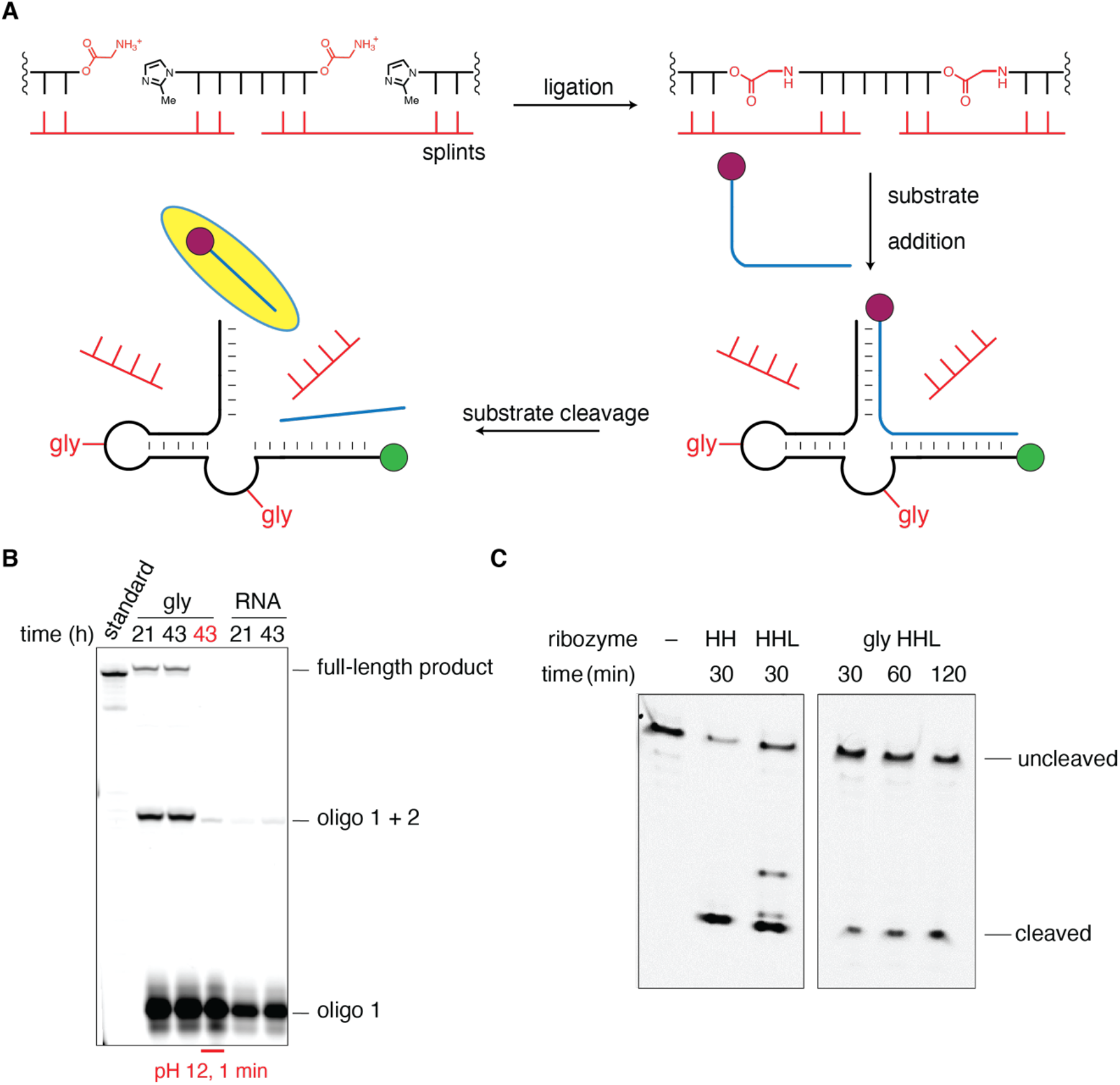
Assembly of a chimeric hammerhead and substrate cleavage in a one pot reaction. **A** Diagram of the splint-assisted assembly method. Oligonucleotides were activated and glycylated as in Figure 2. Splints (red) were 10-nt long such that they made 5 base pairs with each ligating oligonucleotide. The substrate was 5′ labeled with Cy5 (burgundy circle) to easily distinguish the cleaved (highlighted) and uncleaved bands from the FAM labeled hammerhead oligonucleotides. **B** Urea-PAGE of a DNA splint-assisted assembly reaction. Treatment of the glycylated assembly reaction with 200 mM NaOH for 1 minute resulted in the disappearance of the full-length product band, indicating that the product was gly bridged.**C** One-pot hammerhead cleavage reaction at 25 °C. Lanes HH and HHL show substrate cleavage by the all-RNA control ribozymes. HH ribozyme is the identical ribozyme used in Figure 2. HHL is a modified ribozyme used in splint-assisted assembly (see Figure S9). Lanes labeled gly HHL show time-dependent one-pot substrate cleavage by the chimeric gly bridged HHL ribozyme assembled on DNA splints.

## Discussion

Our finding that aminoacylated RNA oligonucleotides can rapidly self-assemble into amino acid bridged RNAs with catalytic activity suggests a possible role for aminoacylation chemistry prior to the evolution of ribosomal translation. Specifically, we outline a scenario for the transition from inefficient non-enzymatic to ribozyme-catalyzed RNA aminoacylation, by showing that a chimeric aminoacylating ribozyme can form from the non-enzymatic assembly of aminoacylated RNAs. On the primordial Earth these processes may have acted in a positive feedback loop, in which a more efficient ribozyme makes more of its components and thus more of itself. This autocatalytic cycle may have driven the early evolution of aminoacyl-RNA synthetase ribozymes. In this context, it is noteworthy that *in vitro* selection and evolution experiments have produced multiple aminoacyl transferase ribozymes with distinct sequences and amino acid substrates, suggesting that ribozyme-catalyzed aminoacylation could have been a function that was relatively easy to evolve in the RNA World.(33–38)

The reduced activity of our chimeric ribozymes compared to their all-RNA progenitors is not surprising considering that these ribozymes were initially evolved from libraries consisting entirely of RNA. The extent of the decrease in catalytic activity depends on the particular ribozyme, the identity of the amino acid, and the sites of the amino acid bridges. The decreased efficiencies could be a direct consequence of local structural distortions or misfolding due to the presence of amino acids interrupting the RNA backbone. However, we speculate that ribozymes that evolved directly through the assembly of aminoacylated RNAs might exhibit more robust activity, as the RNA sequence and the position of the amino acid bridges would be selected to minimize deleterious effects of the bridges. Indeed, the amino acid bridges in chimeric ribozymes could potentially enable the evolution of chimeric ribozymes with enhanced substrate binding or catalytic activity or even activities inaccessible to RNA. Amino acid side chains in the chimeric backbone introduce diverse functional groups not found in RNA and thereby expand the catalytic capabilities of ribozymes. Charged side chains close to a ribozyme active site could promote charge stabilization in a reaction transition state, while side chains with a p*Ka* close to neutral pH, such as the imidazole of histidine, could participate in general acid-base catalysis. Just as the presence of amino acid side chains could allow RNA to perform hitherto inaccessible functions, base pairing could enable chimeric polymers to perform functions that are outside the scope of proteins. Aptamers and ribozymes containing modified nucleobases have been evolved to perform novel functions,(39–46) implying that adding chemical complexity to RNA can be beneficial. The evolution of ribozyme aminoacyl RNA synthetases with distinct amino acid and RNA sequence specificities would have allowed for the assembly of chimeric ribozymes in which specific amino acid bridges were placed consistently at the optimal positions for enhanced chimeric ribozyme activity.

While amino acid bridges in RNA have clear potential advantages, they may also come with certain liabilities. The greater lability of amino acid ester-phosphoramidate linkages to hydrolysis(24, 27) is likely to be responsible for the observed decrease in chimeric ribozyme activity with time. We suggest that the disassembly of a ribozyme into reusable oligonucleotide components could provide additional opportunities for ribozyme regulation. Whereas the facile (dis)assembly of chimeric ribozymes results in a dynamic system, a stable polymer is needed to preserve genetic information. Our assembly of a functional chimeric RNA ligase ribozyme demonstrates that dynamic amino acid-bridged RNAs could also lead to the enzymatic assembly of pure, and therefore more hydrolytically stable genomic RNAs.

The amino acid bridged RNAs reported here unite amino acids and ribonucleotides into a single functional entity thereby serving as a conceptual intermediate between the RNA and protein worlds. In addition to their implications in the origin of life, chimeric ribozymes constitute a novel class of catalytic biopolymers that may lead to opportunities for the *in vitro* evolution of new enzymatic activities by exploiting the diverse chemistries of different amino acid side chains. Our work implicates RNA aminoacylation chemistry in the non-enzymatic assembly of ribozymes, a process that is completely independent of ribosomal protein synthesis. By facilitating the assembly of chimeric ribozymes, aminoacylation could have played a central role in the emergence of RNA-based catalysts, prior to its modern role in the synthesis of protein catalysts.

## Materials and Methods

### General information

All reagents were purchased from Sigma-Aldrich (St. Louis, MO) unless otherwise specified. TurboDNase was purchased from Thermo Scientific (Waltham, MA). Flexizyme “dFx” and the corresponding mutants M1 and M2 were prepared as described elsewhere(47). PCR was performed with Hot Start Taq 2X Master Mix and in-vitro transcription with HiScribe™ T7 Quick High Yield RNA Synthesis Kit from New England Biolabs (Ipswich, MA). EDTA is used as an abbreviation for Na_2_EDTA pH 8.0.

### Oligonucleotide synthesis

Oligonucleotides were either purchased from Integrated DNA Technologies (Coralville, IA) or synthesized in house on an Expedite 8909 solid-phase oligonucleotide synthesizer. Phosphoramidites and reagents for the Expedite synthesizer were purchased from either Glen Research (Sterling, VA) or Chemgenes (Wilmington, MA). Cleavage of synthesized oligonucleotides from the solid support was performed using 1 mL of AMA (1:1 mixture of 28 % aqueous ammonium hydroxide and 40 % aqueous methylamine) for 30 minutes at room temperature, while deprotection was done in the same solution for 20 minutes at 65 °C. Deprotected oligonucleotides were lyophilized, resuspended in 100 μL DMSO and 125 μL TEA.3HF, and heated at 65 °C for 2.5 h to remove TBDMS from 2′ hydroxyls. Following this deprotection, oligonucleotides were purified by preparative 20% polyacrylamide gel electrophoresis (19:1 with 7 M urea), desalted using Waters Sep-Pak C18 cartridges (Milford, MA), and characterized by high-resolution mass spectrometry on an Agilent 6230 TOF mass spectrometer.

### Oligonucleotide activation

Oligonucleotides phosphorylated at the 5′-OH were activated with 2-methylimidazole as previously reported(13) with the following modifications: gel-purified products of 1 μmol solid-phase synthesis were dissolved in 100 μL DMSO. 0.05 mmol of triethylamine (TEA), 0.02 mmol of triphenylphosphine (TPP), 0.04 mmol of 2-methylimidazole, and 0.02 mmol of 2,2′-dipyridyldisulfide (DPDS) were added to the reaction, and the reaction was incubated on a rotator for 5 h at room temperature. After 5 h, all of the reagents above were added in listed quantities again and the reaction was allowed to rotate for an additional 12 h at room temperature. The reaction was precipitated with 100 μL saturated NaClO4 in acetone and 1 mL acetone for 30 minutes on dry ice. The pellet was washed with 1 mL 1:1 acetone:diethylether twice. The products were resolved and purified by HPLC on an Agilent ZORBAX analytical column (Eclipse Plus C18, 250 × 4.6mm, 5 μm particle size, P.N. 959990-902), at a flow rate of 1 ml/min. Gradient: (A) aqueous 20 mM triethylammonium bicarbonate pH 8.0, and (B) acetonitrile, from 7% to 12% B over 12 minutes.

2-aminoimidazole (2AI)-activated substrates for ligase ribozyme reactions were generated by reacting corresponding 5′-monophosphorylated oligonucleotides with 0.2 M 1-ethyl-3-(3 dimethylaminopropyl) carbodiimide (HCl salt) and 0.6 M 2-aminoimidazole (HCl salt, pH adjusted to 6) at room temperature for 2 h. The reaction was desalted by exchanging the reaction buffer with water in Amicon Ultra-0.5 mL 3K centrifugal filters and purified by HPLC as above.

### Oligonucleotide aminoacylation

3,5-dinitrobenzyl esters of amino acids (aa-DBEs), synthesized as described previously(27, 47) were dissolved in neat DMSO and added to aminoacylation reactions at 0 °C. Each aminoacylation reaction contained 50 mM Na^+^-HEPES pH 8.0, 10 mM MgCl_2_, 10 μM oligonucleotide, 10 μM Flexizyme, and 5 mM aa-DBE (20% DMSO final). Reactions were allowed to proceed for 16-18 hours at 0 °C before being used in subsequent steps. To quantify the aminoacylation yields, 1 μL aliquots were quenched in acidic gel buffer (10 mM EDTA, 100 mM NaOAc pH 5.0, 150 mM HCl, 70% v/v formamide) and resolved by 20% acidic polyacrylamide gel (19:1 with 7 M urea, 0.1 M NaOAc pH 5.0). The gel was run for 2 h at 300 V and 4 °C and visualized on a Typhoon 9410 imager. A typical aminoacylation reaction yielded 20-60 % product, measured by band densities in ImageQuant TL 8.1 software. Gels with unlabeled oligonucleotides were stained with SYBR Gold, according to the manufacturer’s instructions, and band intensities were quantified as above.

### Aminoacylated oligonucleotide purification

Oligonucleotides used in the assembly of the chimeric hammerhead ribozyme were used without purification, because gel purification of the full-length chimeric hammerhead was sufficient to remove excess Flexizyme. Full-length chimeric ligase could not be resolved from the excess Flexizyme, and the aminoacylated oligonucleotides were purified away from the Flexizyme by acidic preparative gel electrophoresis prior to the assembly reaction. Similarly, aminoacylated oligonucleotides that were used to generate the chimeric FFlexizyme were gel purified prior to the assembly reaction. Oligonucleotide bands were visualized by UV shadowing, the bands of interest were cut, crushed, and tumbled in 1 mL of 50 mM NaOAc, 5 mM EDTA, pH 5.0 for 3 hours at 4 °C. The extracted oligonucleotides were filtered, concentrated, and desalted using Amicon Ultra-0.5 mL 3K centrifugal filters. 1 μL aliquots of the purified aminoacylated oligonucleotides were analyzed by acidic gel electrophoresis to quantify the aminoacylation yields. Due to the hydrolysis of 2-methylimidazole under acidic conditions, additional bands were present in the gel. The identities of oligonucleotides and relative percentages in each mixture were confirmed by LC-MS.

### Hammerhead (HH) ribozyme assembly (Figures 1, 2, S1-3)

Aminoacylation reactions on a 1 mL scale were performed on oligonucleotides 1 and 2 using gly-DBE, L-lys-DBE, L-leu-DBE, and L-asp-DBE as described above. After the 18 h incubation at 0 °C, each reaction was precipitated by the addition of 100 μL 3 M NaOAc pH 5.5 and 3.9 mL 100% ethanol for 20 minutes on dry ice. Pellets were washed twice with 80% ethanol, dried under vacuum, and resuspended in 100 μL of nuclease-free water. Accurate concentration of the mixture of aminoacylated and non-aminoacylated oligonucleotides could not be determined due to the presence of Flexizyme, and were assumed to be 100 μM. For the RNA control reaction, an identical procedure was followed, except no Flexizyme was used during the aminoacylation reaction. Oligonucleotides prepared in this way were used in the assembly reactions described below.

#### Assembly on an RNA template (Figure 1)

Reactions were set up in triplicate at room temperature by mixing 1.25 μM of oligonucleotides 1-3, 1.25 μM of HH RNA template, 2.5 mM MgCl_2_, and 200 mM Na^+^-HEPES pH 8.0 in a final volume of 60 μL. Reaction aliquots (1 μL) were quenched in 14 μL of quench buffer (50 mM EDTA and 9 μM reverse complement of the template in 90% v/v formamide) at the indicated time points, heated at 92 °C for 2 minutes, and resolved by 20 % polyacrylamide gel electrophoresis (19:1 with 7 M urea). The ligated full-length product was quantified for each time point by integrating and normalizing band intensity in each gel lane the in ImageQuant TL 8.1 software. For the glycylated reaction, the fraction of the full-length product was estimated from the fraction of glycylation quantified by acidic gel electrophoresis as described above. Because the percentage of glycylated oligonucleotides 1 and 2 was 21 and 20%, respectively, the raw full-length product yield was divided by 0.042 to obtain the adjusted product yield.

#### Assembly on a DNA template (Figure 2, S3)

500 μL reactions were set up at room temperature by mixing 20 μM of oligonucleotides 1-3, 20 μM of HH DNA template, 10 mM EDTA, and 200 mM Na^+^-HEPES pH 8.0. After 3 h, the reactions were concentrated using Amicon Ultra-0.5 mL 3K centrifugal filters down to 50 μL. 100 U of TurboDNase and 1× TurboDNase buffer were added to the concentrated reactions to a final volume of 500 μL and incubated for 15 minutes at 37 °C. 1 μL aliquots of the digested reactions were quenched in 14 μL quenching buffer (50 mM EDTA in 90% v/v formamide), and the full-length products were resolved and quantified as above. The digested reactions were concentrated using Amicon Ultra-0.5 mL 3K centrifugal filters down to 50 μL, diluted with 60 μL of 10 mM EDTA in 95% v/v formamide, purified by preparative 20% polyacrylamide gel electrophoresis (19:1 with 7 M urea), extracted in 1 mL of 50 mM NaOAc, 5 mM EDTA, pH 5.0 for 3 h at 4 °C, and desalted with the Zymo Clean and Concentrator Kit. The concentration of each chimeric hammerhead was determined in triplicates by serially diluting a known concentration of the all-RNA standard, resolving the dilutions by denaturing gel electrophoresis, quantifying the band intensities, and generating a standard curve.

#### Assembly on splint oligonucleotides (Figures 5 and 9)

30 μL reactions were set up by mixing 25 μM of oligonucleotides 1-3, 25 μM of splints (either DNA or RNA), 100 mM NaCl, 5 mM MgCl_2_, and 200 mM Na^+^-HEPES pH 8.0 at 0 °C. Reaction aliquots (0.5 μL) were quenched in 19.5 μL of quenching buffer (50 mM EDTA in 90% v/v formamide) at the indicated time points, heated at 92 °C for 1 minute, and resolved by 20 % polyacrylamide gel electrophoresis (19:1 with 7 M urea). After 43 h, the assembly reactions were directly used in the activity assay described below.

### Hammerhead ribozyme activity assay (Figures 2, 5, S9)

Hammerhead cleavage was assayed in 10 μL reactions which contained 0.12 μM ribozyme, 0.1 μM FAM/Cy5-labeled substrates, 100 mM Tris-Cl (pH 8), and 3 mM MgCl_2_. To test cleavage activity of the chimeric hammerhead ribozyme assembled on DNA or RNA splints, the 3′ end of the ribozyme was extended by 6 nucleotides, and the corresponding 5′ end of the substrate was extended by 4 nucleotides, resulting in an 11 base pair stem in the ribozyme-substrate complex instead of the usual 5 base paired stem. This was done to favor the formation of a catalytically active hammerhead ribozyme-substrate complex over the inactive ribozyme-splint complex resulting from aminoacylated oligonucleotide ligation. To facilitate the separation of the chimeric ribozyme product from splint oligonucleotides, the ribozyme-splint complex was heated at 95 °C for 2 minutes followed by incubation at either 25 °C or 42 °C for 5 min in the presence of 100 mM Tris-Cl (pH 8) and 0.1 μM Cy5-labeled substrate. Cleavage was initiated by adding 3 mM MgCl_2_ and the reactions were incubated at either 25 °C or 42 °C. 1.5 μL aliquots were quenched in 5 μL loading buffer (7 M urea, 100 mM EDTA in 1X TBE) at various times points and stored on dry ice. Cleaved products were separated from uncleaved substrates by 20% polyacrylamide gel electrophoresis (19:1 with 7 M urea). Bands were quantified with ImageQuant IQTL 8.1 software. Kinetic plots were nonlinearly fitted to the modified first order rate equation, y = A (1 – e^−*k*x^), where A represents the fraction of active complex, k is the first order rate constant, x is time, and y is the fraction of cleaved product in GraphPad Prism 9.

### 2AI-ligase assembly (Figures 3, S4-6)

Aminoacylation reactions on a 4 mL scale were performed on oligonucleotides 1-3 with L-lys-DBE and gly-DBE as described above. After the 18 h incubation at 0 °C, each reaction was precipitated by the addition of 400 μL 3 M NaOAc pH 5.5 and 15.6 mL 100% ethanol for 20 minutes on dry ice. Pellets were washed twice with 80% ethanol, dried under vacuum, and dissolved in 100 μL of 10 mM EDTA in 95% v/v formamide. The oligonucleotides were then purified using preparative acidic gel electrophoresis as described above.

The assembly reaction was set up by mixing 16.7 μM of oligonucleotides 1-4, 16.7 μM ligase RNA-DNA hybrid template, 5 mM EDTA, and 100 mM NaOAc pH 5.0 in a final volume of 300 μL. The reaction was annealed by heating it to 70 °C for 2 minutes and slowly cooling to 4 °C at the rate of 0.1 °C/s. The annealed reaction was diluted to 500 μL with Na^+^-HEPES pH 8.0 at room temperature (final concentrations: 10 μM oligonucleotides 1-4, 10 μM RNA-DNA hybrid template, 3 mM EDTA, 60 mM NaOAc, 200 mM Na^+^-HEPES pH 8.0). The reaction was allowed to proceed for 3 hours before being concentrated, digested, purified, and desalted as described for the hammerhead ribozyme assembly. Concentrations were determined as in the case of the chimeric hammerhead, except that an all-RNA ligase standard was used to generate the standard concentration curve.

### 2AI-ligase ribozyme activity assay (Figures 3, S6)

AI-ligase activity was assayed in 5 μL reactions which contained 1 μM ribozyme, 1.2 μM RNA template, 2 μM 2-aminoimidazole (2AI) 100 mM Tris-Cl (pH 8), 250 mM NaCl, and 10 mM MgCl_2_. 1 μL aliquots were quenched in 5 μL loading buffer (7 M urea, 100 mM EDTA in 1X TBE) at various times points and stored on dry ice. Ligated products were separated from unligated precursors by 10% polyacrylamide gel electrophoresis (19:1 with 7 M urea). Bands were quantified with ImageQuant TL 8.1 software. Kinetic plots were nonlinearly fitted to the modified first order rate equation, y = A (1 − e^−*k*x^), where A represents the fraction of active complex, *k* is the first order rate constant, x is time, and y is the fraction of cleaved product in GraphPad Prism 9.

### Flexizyme assembly (Figures 4, S7-8)

Aminoacylation reactions on a 4 mL scale with L-lys-DBE and gly-DBE were performed and purified as described in the ligase assembly section.

The assembly reaction was set up at room temperature by mixing 15.4 μM of oligonucleotides 1-3, 15.4 μM Flexi DNA template, 3 mM EDTA, and 200 mM Na^+^-HEPES pH 8.0 in a final volume of 500 μL. The reaction was allowed to proceed for 3 hours, before being concentrated, digested, and purified, as described above. Concentrations were determined as for the chimeric hammerhead, except that an all-RNA Flexizyme standard was used to generate the standard concentration curve.

### Flexizyme activity assay (Figures 4, S8)

Flexizyme activity was assayed as follows. 2 μL of 7.5 μM chimeric or all-RNA Flexizyme was treated with 200 mM NaOH for 30 seconds at room temperature before being quenched with equimolar HCl (final volume of the quenched reactions is 3 μL). In parallel, 7.5 μM chimeric or all-RNA Flexizyme was diluted with water to 5 μM and incubated for 30 seconds at room temperature in triplicate. The reactions were diluted to 2.5 μM Flexizyme with 50 mM Na^+^-HEPES pH 8.0, 5 mM gly-DBE (20% DMSO), 10 μM substrate oligonucleotide, 10 mM MgCl_2_, and incubated at 0 °C. Reaction aliquots (0.5 μL) were quenched in 4.5 μL acidic quenching buffer (10 mM EDTA, 100 mM NaOAc pH 5.0, 150 mM HCl, 70 % v/v formamide) at the indicated time points and resolved by 20% acidic polyacrylamide gel (19:1 with 7 M urea, 0.1 M NaOAc pH 5.0). The gel was run for 2 h at 300 V and 4 °C and visualized with a Typhoon 9410 imager. Aminoacylation percentages were quantified by measuring band densities in ImageQuant TL 8.1 software. The remaining aminoacylation reactions at 22 h were desalted using Zip-Tip C18 columns and characterized by high-resolution mass spectrometry on an Agilent 6230 TOF mass spectrometer.

## Acknowledgments

The authors acknowledge Drs. Daniel Duzdevich, Lijun Zhou, Long-Fei Wu, Victor S. Lelyveld, and Seohyun Chris Kim for their helpful suggestions.

