## Supplementary material for "Non-enzymatic assembly of active chimeric ribozymes from aminoacylated RNA oligonucleotides": SI

[a]Howard Hughes Medical Institute, Department of Molecular Biology and Center for Computational and Integrative Biology, Massachusetts General Hospital, 185 Cambridge Street, Boston, Massachusetts 02114, United States; [b]Department of Genetics, Harvard Medical School, Boston, Massachusetts 02115, United States; [c]Department of Chemistry and Chemical Biology, Harvard University, Cambridge, Massachusetts 02138, United States.

[1]These authors contributed equally

**This PDF file includes:**

Figures S1 to S9
Table S1

### Supplementary Figures

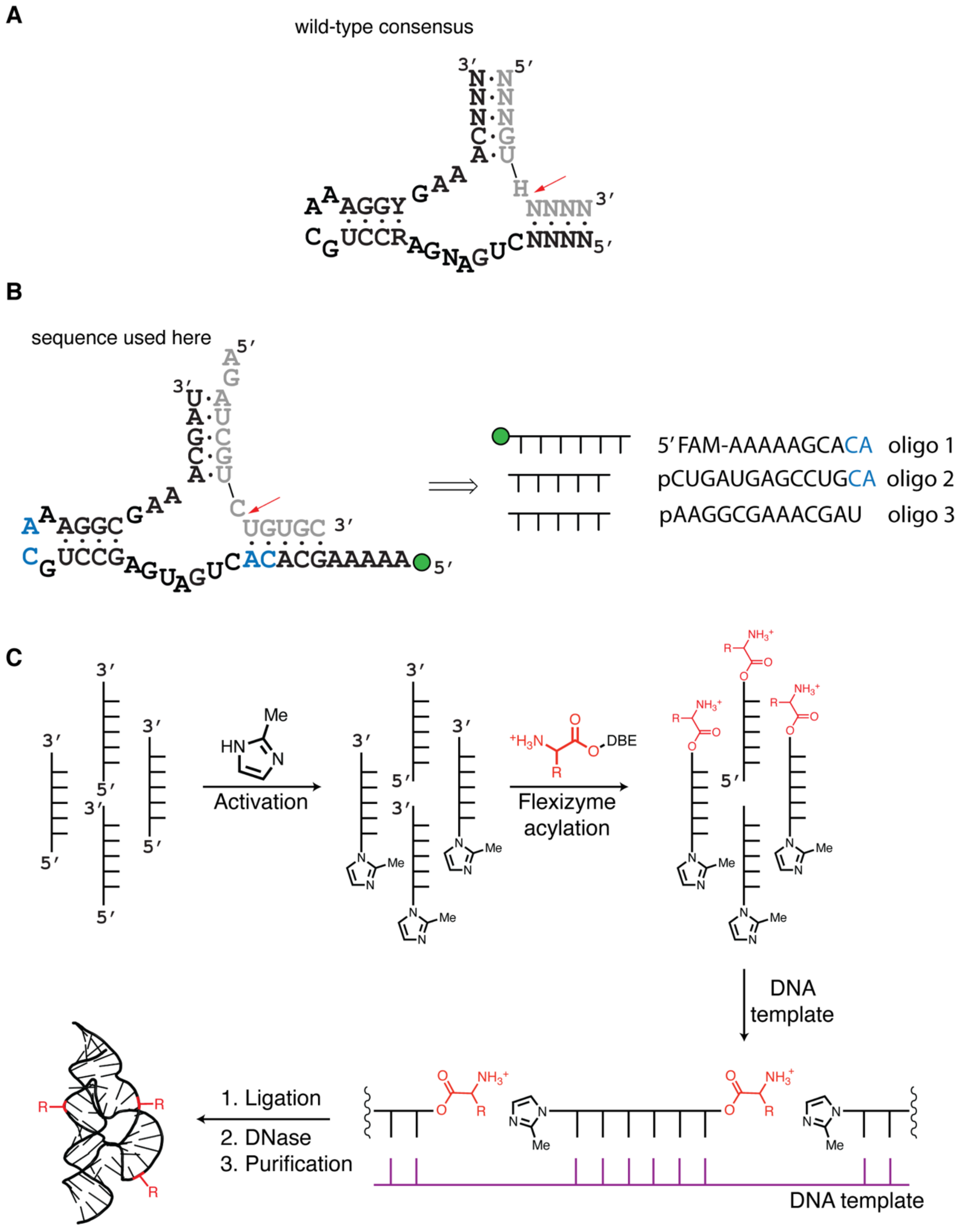

**Figure S1. Hammerhead mutation and deconstruction. A** The wild-type consensus sequence of the hammerhead ribozyme. **B** Sequence variant used in this work to accommodate flexizyme-catalyzed aminoacylation. The hammerhead variant sequence was assembled from three oligonucleotides of roughly equal length (designated oligos 1-3). Blue nucleotides represent the 5′-CA-3′ dinucleotides that are the required substrates for the flexizyme-catalyzed aminoacylation. 5′ $A_5$ sequence was added to increase aminoacylation yield. Red arrow represents the cleavage site in the RNA substrate. The green circle represents the FAM label. The "p" prefix represents the 5′ phosphate. **C** Diagram of the method used to assemble chimeric ribozymes. Oligonucleotides that comprise the ribozymes are first activated with 2-methylimidazole. The oligonucleotides, except for the 3′ terminal oligonucleotide, are then aminoacylated using the Flexizyme ribozyme and 2,4-dinitrobenzyl esters of amino acids. Following the addition of a DNA template, ligation occurs, and subsequent digestion of the template with DNase allows isolation of the ribozyme by preparative urea-PAGE.

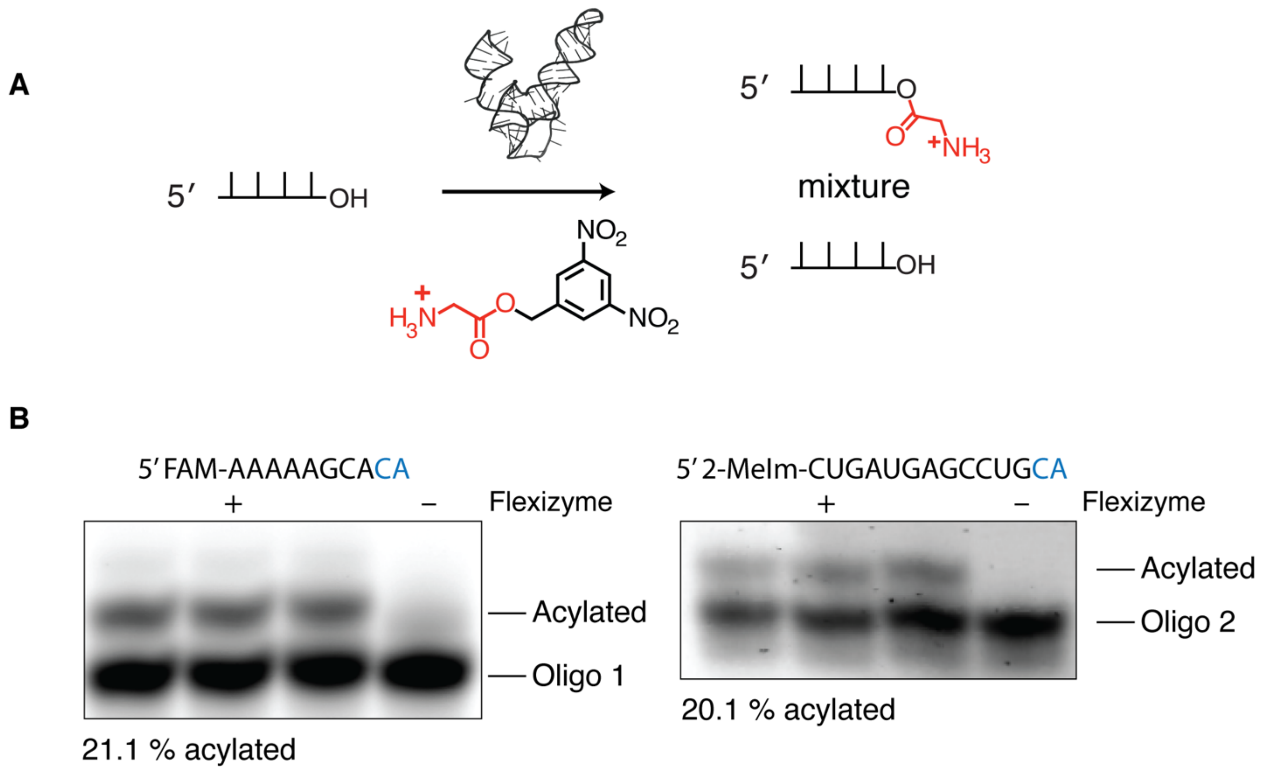


**Figure S2. Aminoacylation of RNA oligonucleotides that comprise the hammerhead ribozyme. A** Schematic of the Flexizyme-catalyzed aminoacylation reaction. Flexizyme accepts 3,5-dinitrobenzyl esters of amino acids as substrates and specifically aminoacylates the 3′ OH of the *cis* diol of RNA oligonucleotides. The reaction does not proceed to completion and thus yields a mixture of aminoacylated and non-aminoacylated RNA. **B** Aminoacylation of oligos 1 and 2 was monitored by 20% denaturing acid urea-PAGE (0.1 M NaOAc pH 5.0, 7 M urea; running buffer 0.1 M sodium acetate pH 5.0). The aminoacylated oligonucleotide migrates slower than the non-aminoacylated oligonucleotide and full resolution of the two bands allows quantification of the aminoacylation yield. Oligo 1 was fluorescently labeled, thereby allowing direct quantification, whereas oligo 2 was stained with SYBR Gold for quantification. The yields below each gel represent an average of triplicate measurements.

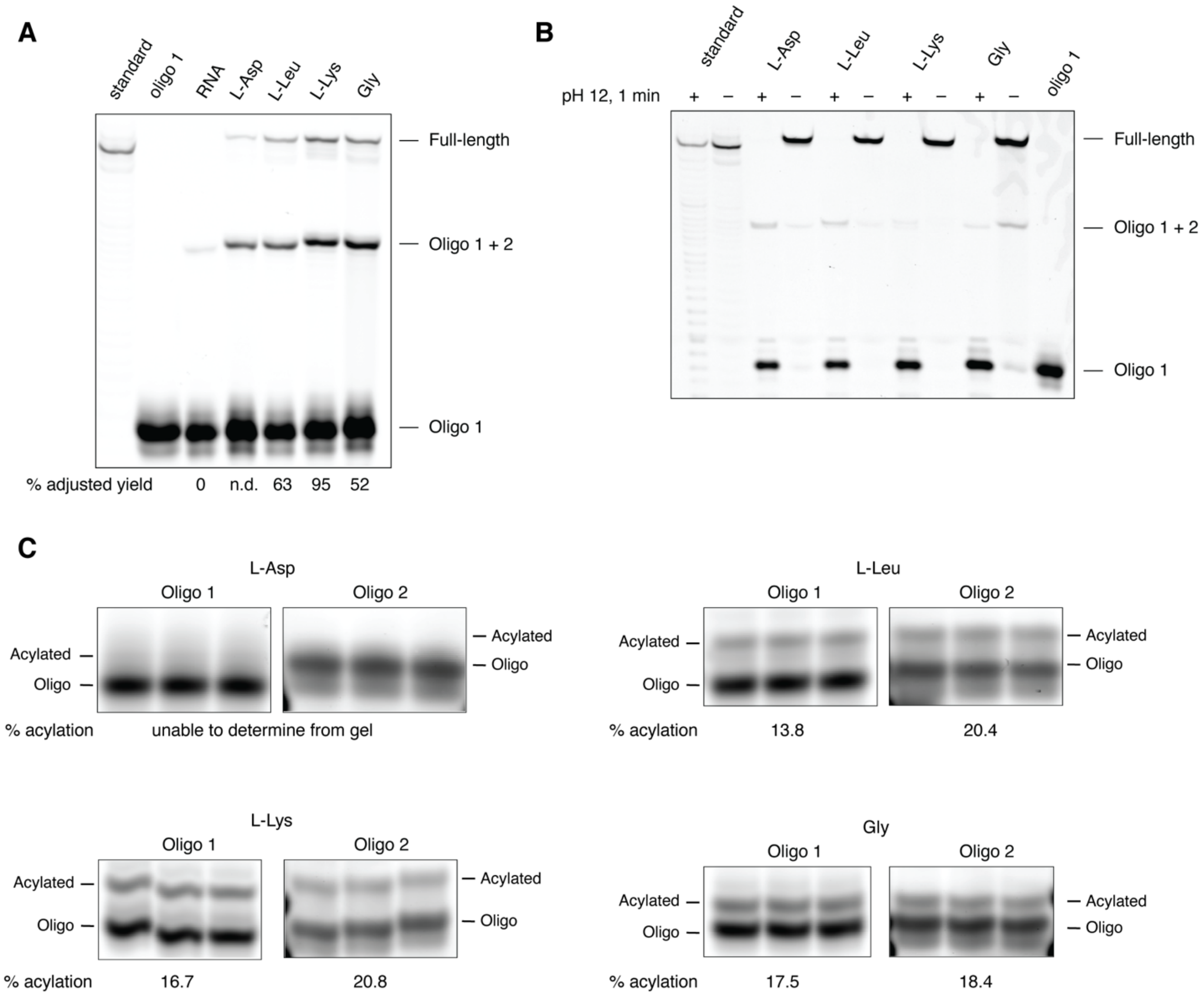


**Figure S3. Chimeric hammerhead assembly with four different amino acids. A** A representative denaturing 20% urea-PAGE gel of the hammerhead assembly reactions. The true yield of the L-Asp assembly reaction could not be determined due to the poor resolution of the aminoacylated and non-aminoacylated bands by acidic urea-PAGE. The standard was a 5′ FAM-labeled hammerhead ribozyme sequence purchased from IDT. **B** Each purified, chimeric ribozyme was subjected to transient alkaline conditions by the addition of 200 mM NaOH for 1 minute. After the NaOH treatment, all four chimeric ribozymes were hydrolyzed such that no full-length product was detectable. The RNA standard displayed minor non-specific hydrolysis. A faint band that corresponds to “Oligo 1 + 2” is still visible due to the incomplete hydrolysis and/or background RNA reaction during assembly. **C** Acidic urea-PAGE analysis of aminoacylation reactions for the four different amino acids. Percent acylation represents the average of technical triplicates. These percent values were used to normalize the assembly yield as described in Methods. L-Asp aminoacylated RNA could not be resolved from non-aminoacylated RNA, hence the adjusted yield for this assembly could not be calculated.

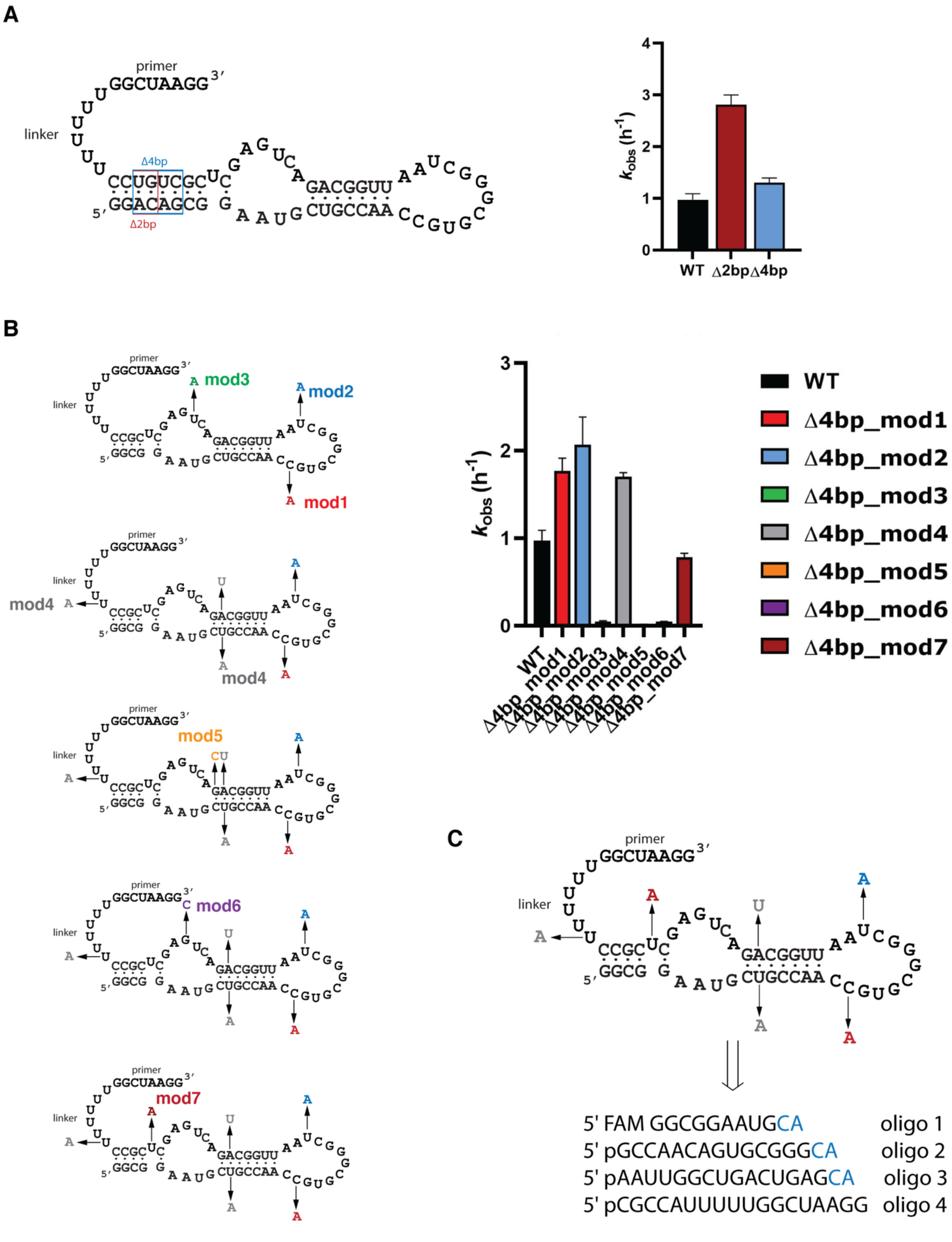


**Figure S4. Optimization of 2AI ligase ribozyme sequence. A** Truncations of the base pairs within the blue and red rectangles yielded shorter ribozymes that retained the catalytic RNA ligation activity. **B** The 4 bp deletion mutant from **A** was altered to install the 5′-CA-3′ dinucleotide required for Flexizyme substrates. The sequence labeled “mod7” displayed similar catalytic RNA ligation activity to the 4 bp deletion mutant and was selected for the assembly experiments. **C** The “mod7” ligase ribozyme was assembled from four oligonucleotides ranging from 11 nt to 18 nt in length (designated oligos 1-4). Sequences labeled in blue represent the aminoacylation sites. The “p” prefix represents the 5′ phosphate.

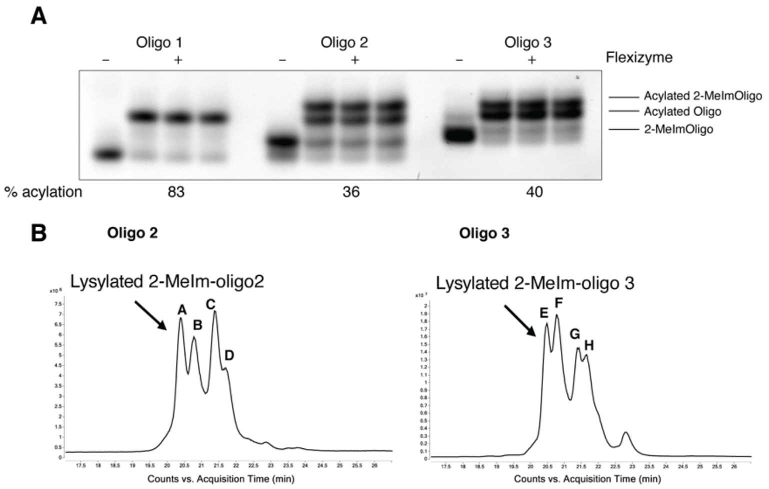


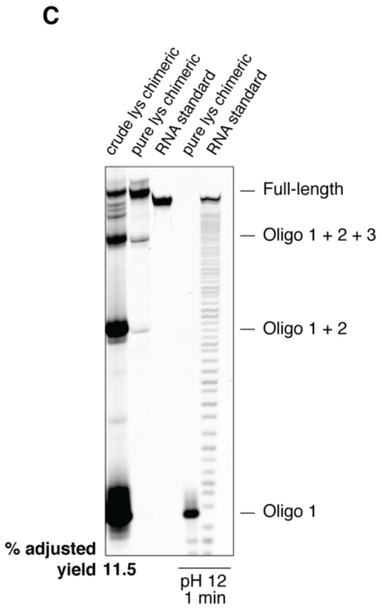


| Signal | Calculated | Found | Identity |
|---|---|---|---|
| A | 5427.8745 | 5427.7797 | Lysylated 2-MeIm-oligo 2 |
| B | 5363.8320 | 5363.7192 | Lysylated oligo 2 |
| C | 5299.7795 | 5299.6727 | 2-MeIm-oligo 2 |
| D | 5235.7370 | 5235.6360 | oligo 2 |
| E | 5719.8729 | 5719.7686 | Lysylated 2-MeIm-oligo 3 |
| F | 5655.8304 | 5655.7277 | Lysylated oligo 3 |
| G | 5591.7779 | 5591.6872 | 2-MeIm-oligo 3 |
| H | 5527.7354 | 5527.6432 | oligo 3 |

**Figure S5. Assembly of the chimeric RNA ligase from lysylated oligonucleotides. A** Acidic urea-PAGE analysis of aminoacylation reactions for the three different oligonucleotides stained with SYBR Gold. The lysylated oligonucleotide migrates more slowly than the non-lysylated oligonucleotide. Additional bands present in oligo 2 and oligo 3 samples represent lysylated unactivated oligonucleotides due to hydrolysis of 2-methylimidazole during acidic gel purification. **B** Top: total ion chromatograms for oligos 2 and 3 that were used to determine the identity of the additional bands shown in **A**. Bottom: UV chromatograms for oligos 2 and 3 that were used to estimate the percentage of lysylated, activated oligonucleotides in each sample. These values roughly matched the gel quantitation values and were used to normalize the assembly yield as described in Methods. **C** A representative denaturing 20% urea-PAGE gel of the L-lys ligase assembly reaction. Assembly reaction was performed as described in the Methods, and the adjusted yield was the average of technical triplicates. The standard was a 5′ FAM-labeled ligase sequence purchased from IDT. The purified chimeric ribozyme was subjected to transient alkaline conditions by the addition of 200 mM NaOH for 1 minute. After the NaOH treatment, the chimeric ribozyme was hydrolyzed such that no full-length product was detectable. The RNA standard displayed minor non-specific hydrolysis.

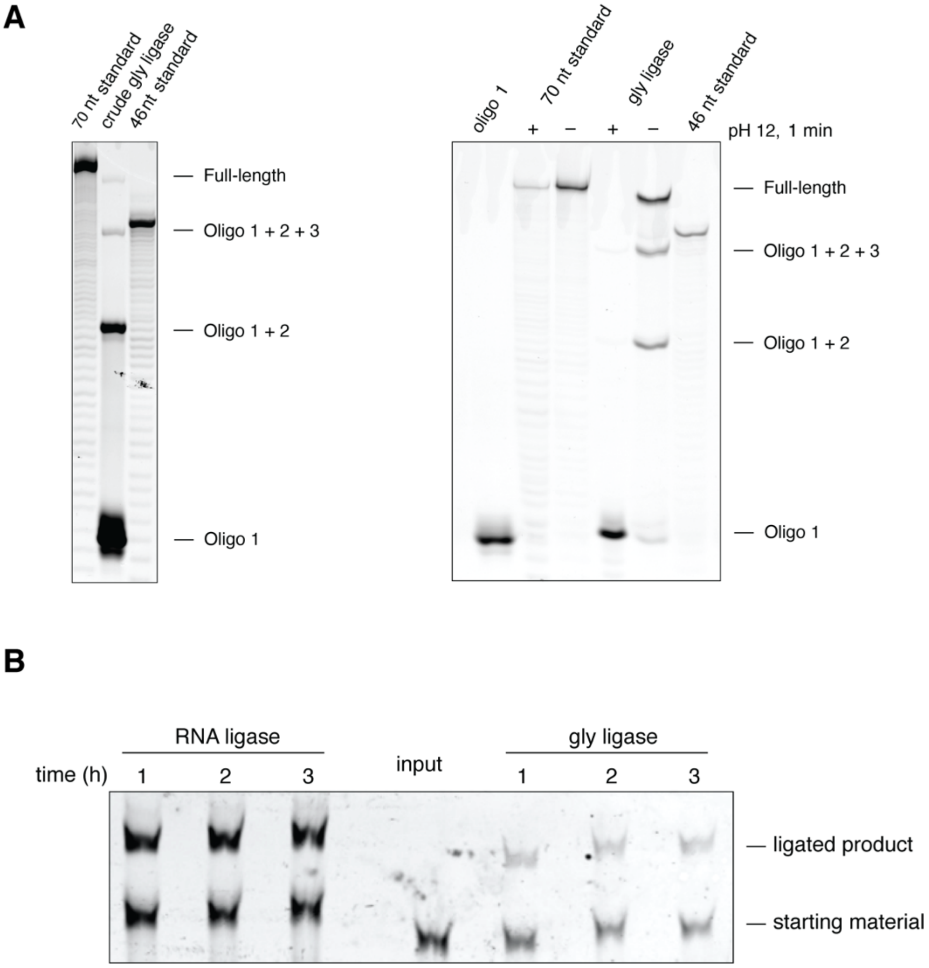


**Figure S6. Assembly of a chimeric RNA ligase from glycylated oligonucleotides. A** Left: A representative denaturing 20% urea-PAGE gel of the gly ligase assembly reaction. Assembly reaction was performed as described in the Methods. The adjusted yield was not calculated. The two standards used were 5′ FAM-labeled 70 nt and 46 nt sequences purchased from IDT. Right: The purified chimeric ribozyme was subjected to transient alkaline conditions by the addition of 200 mM NaOH for 1 minute. After the NaOH treatment, the chimeric ribozyme was hydrolyzed such that no full-length product was detectable. The RNA standards displayed minor non-specific hydrolysis. **B** A representative denaturing 20% urea-PAGE of the gly-ligase ligation reaction over time. The ligation product band is clearly visible; however, due to low ligation yields caused by competing hydrolysis of the phosphoramidate linkages in the product and ribozyme, we did not determine the product yields.

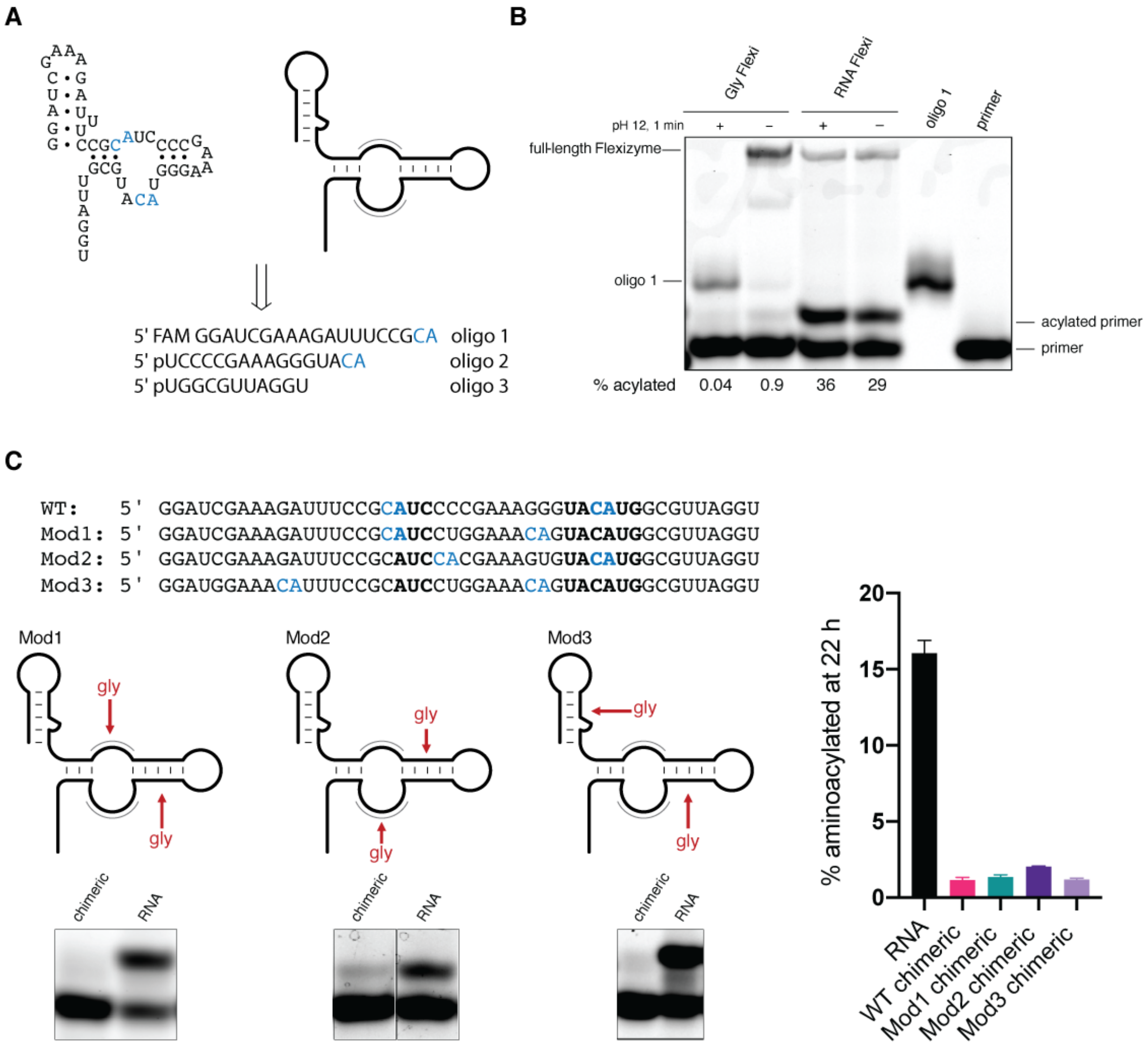


**Figure S7. A functional chimeric Flexizyme can be assembled from glycylated RNA. A** Left: The wild-type Flexizyme sequence; right: diagram of Flexizyme used hereafter. CA sequences that serve as substrates for the Flexizyme aminoacylation are shown in blue. The Flexizyme was assembled from three oligonucleotides (designated oligos 1-3). The “p” prefix represents the 5′ phosphate. **B** A representative denaturing 20% acidic urea-PAGE used to monitor the Flexizyme activity at the 22 h time point. The gly chimeric Flexizyme retained the aminoacylating activity of its all-RNA counterpart, which disappeared after transient alkaline treatment. **C** The wild-type Flexizyme sequence was mutated to reposition the CA sequences and gly bridges outside the catalytic center, which is indicated by a grey outline. Each Flexizyme variant was modestly active as seen from and representative denaturing 20% acidic urea-PAGE. Aminoacylation yields of each ribozyme were determined by averaging triplicate measurements (right).

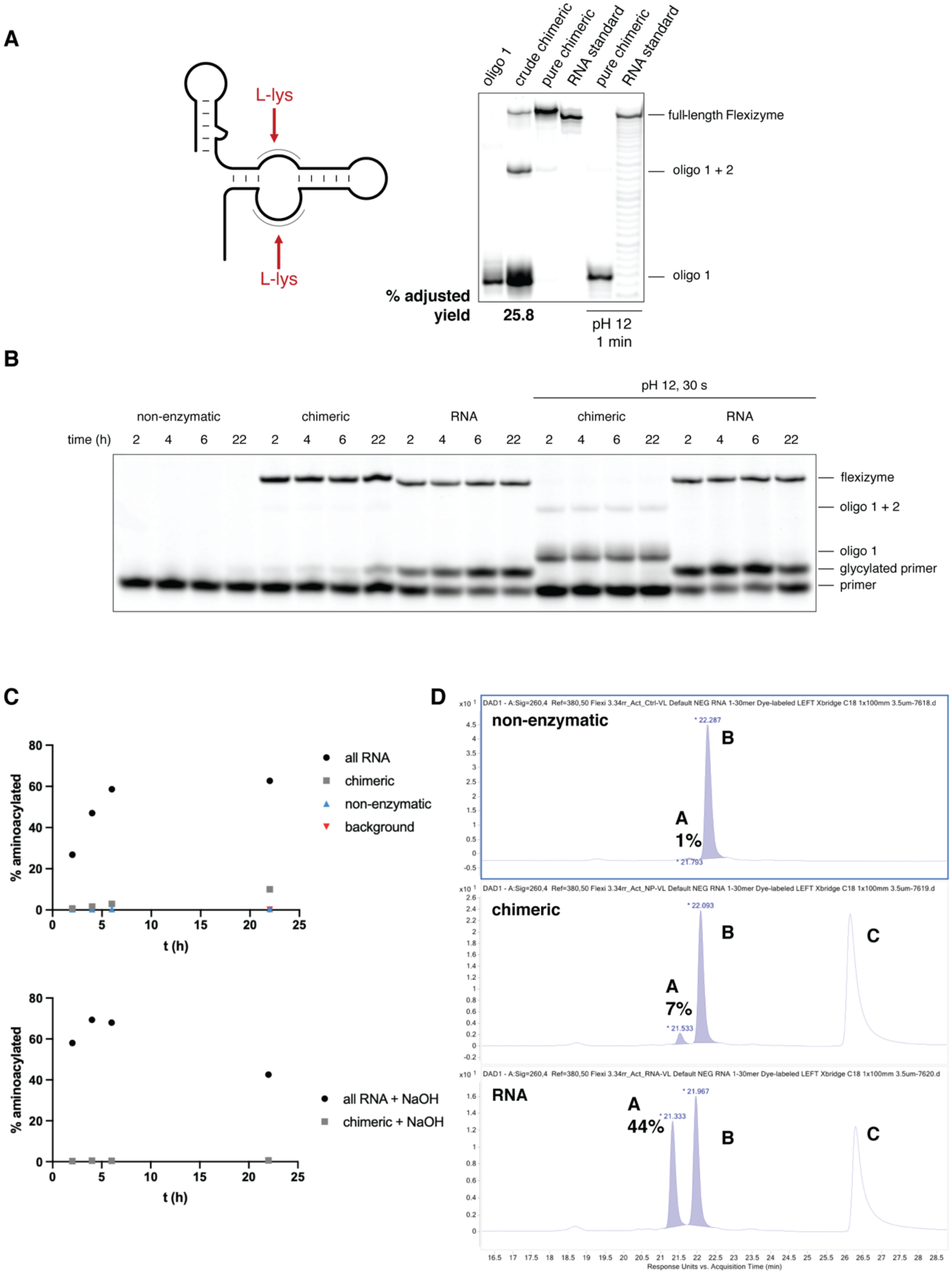


| Signal | Calculated | Found | Identity |
|---|---|---|---|
| A | 3822.6720 | 3822.6312 | glycylated primer |
| B | 3765.6506 | 3765.6234 | primer |
| C | N/A | N/A | flexizyme |

**Figure S8. Chimeric Flexizyme assembled from L-lysylated RNA is functional. A** Left: diagram of the Flexizyme with the L-lys “bridges” in the catalytic center indicated by the red arrows. Right: representative denaturing 20% urea-PAGE of the L-lys Flexizyme assembly reaction. The purified chimeric ribozyme was subjected to transient alkaline conditions by the addition of 200 mM NaOH for 1 minute. After the NaOH treatment, the chimeric ribozyme was hydrolyzed such that no full-length product was detectable. The RNA standard displayed minor non-specific hydrolysis. **B** A representative denaturing 20% acidic urea-PAGE used to monitor the Flexizyme activity over time. The non-enzymatic reaction included all components except for the Flexizyme. The chimeric L-lys Flexizyme aminoacylates the primer substrate with glycine only if it is not pretreated with 200 mM NaOH. The RNA standard Flexizyme retains its activity even after the alkaline pretreatment. **C** Top: the aminoacylation activity was monitored over time in triplicate and the average percent aminoacylation was plotted. The background value was determined by loading the pure primer substrate and quantifying the amount of signal at the gel location parallel to the aminoacylated band. Bottom: aminoacylation activity monitored over time after the pretreatment with 200 mM NaOH for 30 seconds. **D** UV chromatograms of the three reactions collected at the 22 h time point of the aminoacylation reaction. Integrating the signals that corresponded to the aminoacylated primer and primer resulted in values that roughly match the gel analysis. Some hydrolysis of the aminoacylated primer is expected during LC-MS analysis, hence the lower values compared to the gel. The calculated and observed m/z for each UV signal are tabulated, confirming that aminoacylation with glycine occurs.

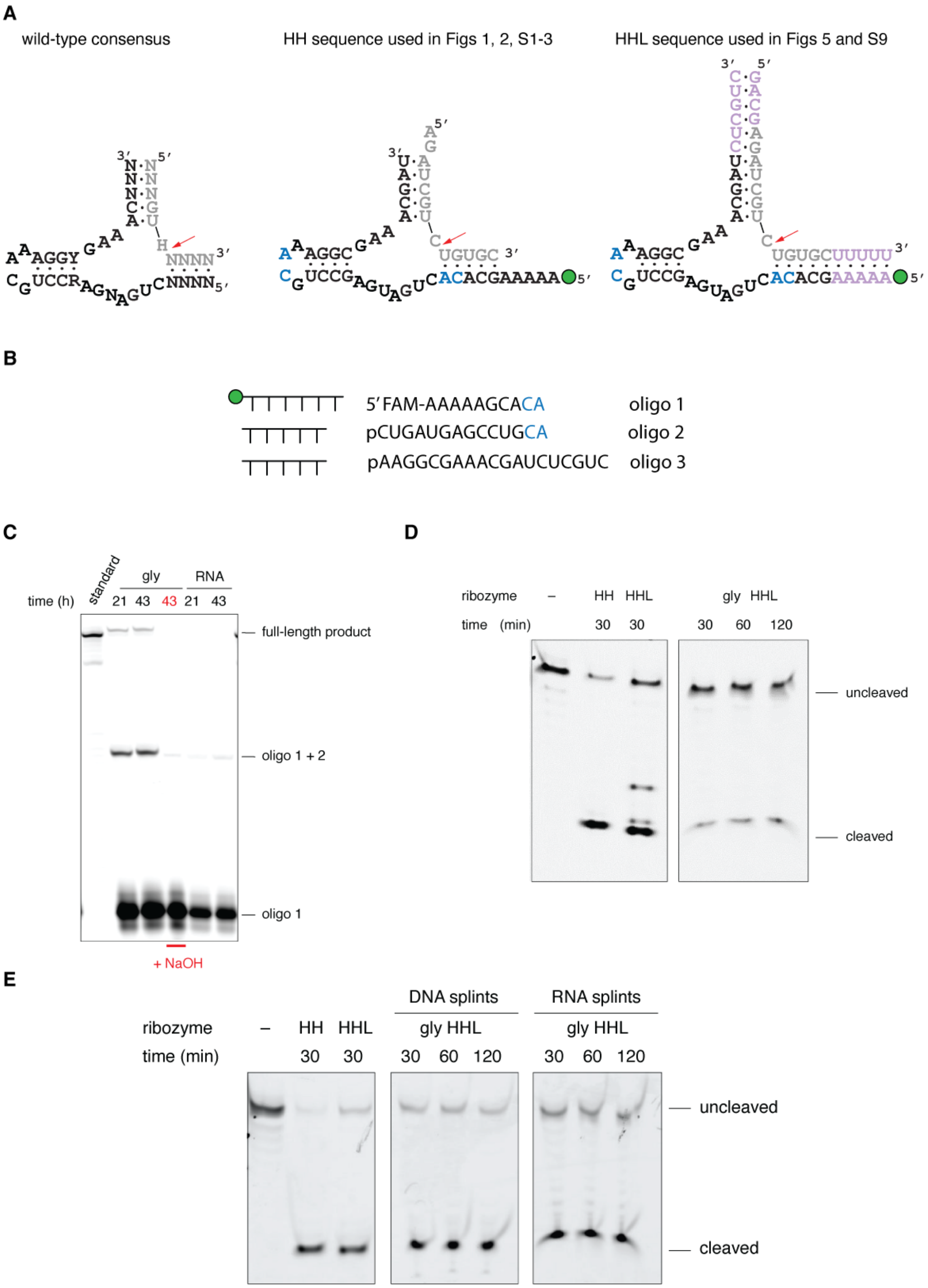


**Figure S9. Chimeric hammerhead assembled using splints from glycylated oligonucleotides cleaves its substrate in the same pot. A** Diagram of the hammerhead ribozymes used in this study. **B** Oligonucleotides used in the splint-assisted assembly of the chimeric HHL ribozyme. **C** Urea-PAGE of an RNA splint-assisted assembly reaction. Treatment of the glycylated assembly reaction with 200 mM NaOH for 1 minute resulted in the disappearance of the full-length product band, indicating that the product was gly bridged. **D** One-pot hammerhead cleavage reaction at 25 ºC. Lanes HH and HHL show substrate cleavage by the all-RNA control ribozymes. Lanes labeled gly HHL show time-dependent one-pot substrate cleavage by the chimeric HHL ribozyme assembled on RNA splints. **E** One-pot hammerhead cleavage reaction at 42 ºC.

**Supplementary Table**

**Table S1:** Sequences used in this work. Nucleotides in purple are deoxyribonucleotides, while all others are ribonucleotides. 2-MeImp represents 2-methylimidazole activated phosphate. 2-AImp represents 2-aminoimidazolde activated phosphate.

| Name | Sequence | Use |
|---|---|---|
| dFx Flexizyme M1 | 5′-GGAUCGAAAGAUUUCCGCAUCCCC GAAAGGGUACAUGGCGUUAGCU | Figures 1-4, S2,3,5-8 |
| dFx Flexizyme M2 | 5′-GGAUCGAAAGAUUUCCGCAUCCCC GAAAGGGUACAUGGCGUUAGUU | Figures 1-4, S2,3,5-8 |
| HH oligo 1 | 5′-FAM-AAAAAGCACA | Figures 1, 2, S1-3 |
| HH oligo 2 | 5′-2-MeImpCUGAUGAGCCUGCA | Figures 1, 2, S1-3 |
| HH oligo 3 | 5′-2-MeImpAAGGCGAAACGAU | Figures 1, 2, S1-3 |
| HH oligo 3L | 5′-2-MeImpAAGGCGAAACGAUCUCGUC | Figures 5, S9 |
| HH RNA template | 5′-AUCGUUUCGCCUUUGCAGGCUCAUCAGUGUGCUUUUU | Figure 1 |
| HH DNA template | 5′-ATCGTTTCGCCTTTGCAGGCTCATCAGTGTGCTTTTT | Figures 2, S3 |
| HH standard | 5′-FAM-AAA AAG CAC ACU GAU GAG CCU GCA AAG GCG AAA CGA U | Figures 1,2, S3 |
| HHL standard | 5′-FAM-AAA AAG CAC ACU GAU GAG CCU GCA AAG GCG AAA CGA UCU CGU C | Figures 5, S9 |
| HH substrate | 5′-FAM-AGAUCGUCUGUGC | Figure 2 |

| | | |
|---|---|---|
| HH DNA splint 1 | 5′-ATCAGTGTGC | Figures 5, S9 |
| HH DNA splint 2 | 5′-GCCTTTGCAG | Figures 5, S9 |
| HH RNA splint 1 | 5′-AUUGGUGUGU | Figure S9 |
| HH RNA splint 2 | 5′-GUUUUUGUAG | Figure S9 |
| HHL substrate | 5′-Cy5-GACGAGAUCGUCUGUGCUUUUU | Figures 5, S9 |
| Ligase oligo 1 | 5′-FAM-GGCGGAAUGCA | Figures 3, S4-6 |
| Ligase oligo 2 | 5′-2-MeImpGCCAACAGUGCGGGCA | Figures 3, S4-6 |
| Ligase oligo 3 | 5′-2-MeImpAAUUGGCUGACUGAGCA | Figures 3, S4-6 |
| Ligase oligo 4 | 5′-2-MeImpCGCCAUUUUUGGCUAAGG | Figures 3, S4-6 |
| Ligase DNA-RNA template | 5′-CCTTAGCCAAAAATGGCGUGCTCAGTCAGCCAAUUUGCCCGCACTGTTGGCUGCATTCCGCC | Figures 3, S5,6 |
| Ligase standard | 5′-FAM-GGCGGAAUGCAGCCAACAGUGCGGGCAA AUUGGCUGACUGAGCACGCCAUUUUUGGCUAAGG | Figures 3, S5 |
| 70-nt standard | 5′-FAM-GGACAGCGGAAUGCUGCCAACCGUGCGGGCUAAUUG GCAGACUGAGCUCGCUGUCCUUUUUGGCUAAGG | Figure S6 |
| 46-nt standard | 5′-FAM-GGAUCGAAAGAUUUCCGCAUCCCC GAAAGGGUACAUGGCGUUAGCU | Figure S6 |
| Ligase substrate | 5′-2-AimpACCACCGCAUUCCGCA | Figures 3, S6 |

| | | |
|---|---|---|
| Flexizyme oligo 1 | 5′-FAM-GGAUCGAAAGAUUUCCGCA | Figures 4, S7-8 |
| Flexizyme oligo 2 | 5′-2-MeImpUCCCCGAAAGGGUACA | Figures 4, S7-8 |
| Flexizyme oligo 3 | 5′-2-MeImpUGGCGUUAGGU | Figures 4, S7-8 |
| Flexizyme DNA template | 5′-ACCTAACGCCATGTACCCTTTCGGGGATGCGGAAATCTTTCGATCC | Figures 4, S7-8 |
| Flexizyme standard | 5′-FAM-GGAUCGAAAGAUUUCCGCAUCCCCGAAAGGGUACAUGGCGUUAGGU | Figures 4, S7-8 |
| Flexizyme mod1 oligo 1 | 5′-FAM-GGAUCGAAAGAUUUCCGCA | Figure S7 |
| Flexizyme mod1 oligo 2 | 5′-2-MeImpUCCUGGAAACA | Figure S7 |
| Flexizyme mod1 oligo 3 | 5′-2-MeImpGUACAUGGCGUUAGGU | Figure S7 |
| Flexizyme mod1 DNA template | 5′-ACCTAACGCCATGTACTGTTTCCAGGATGCGGAAATCTTTCGATCC | Figure S7 |
| Flexizyme mod1 standard | 5′-FAM-GGAUCGAAAGAUUUCCGCAUCCUGGAAACAGUACAUGGCGUUAGGU | Figure S7 |
| Flexizyme mod2 oligo 1 | 5′-FAM-GGAUCGAAAGAUUUCCGCAUCCA | Figure S7 |
| Flexizyme mod2 oligo 2 | 5′-2-MeImpCGAAAGUGUACA | Figure S7 |

| | | |
|---|---|---|
| Flexizyme mod2 oligo 3 | 5′-2-MeImpUGGCGUUAGGU | Figure S7 |
| Flexizyme mod2 DNA template | 5′-ACCTAACGCCATGTACACTTTCGTGGATGCGGAAATCTTTCGATCC | Figure S7 |
| Flexizyme mod2 standard | 5′-FAM-GGAUCGAAAGAUUUCCGCAUCCACGAAAGUGUACAUGGCGUUAGGU | Figure S7 |
| Flexizyme mod3 oligo 1 | 5′-FAM-GGAUGGAAACA | Figure S7 |
| Flexizyme mod3 oligo 2 | 5′-2-MeImpUUUCCGCAUCCUGGAAACA | Figure S7 |
| Flexizyme mod3 oligo 3 | 5′-2-MeImpGUACAUGGCGUUAGGU | Figure S7 |
| Flexizyme mod3 DNA template | 5′-ACCTAACGCCATGTACTGTTTCCAGGATGCGGAAATGTTTCCATCC | Figure S7 |
| Flexizyme mod3 standard | 5′-FAM-GGAUGGAAACAUUUCCGCAUCCUGGAAACAGUACAUGGCGUUAGGU | Figure S7 |
| Flexizyme substrate | 5′-FAM-AGAGAAGCCA | Figures 4, S7, S8 |
